# A Vascularized Liver Microphysiological System Captures Key Features of Hepatic Insulin Resistance and Monocyte Infiltration

**DOI:** 10.1101/2025.01.09.632182

**Authors:** Erin N. Tevonian, Ellen L. Kan, Kairav K. Maniar, Alex J. Wang, Anisha Datta, Roger D. Kamm, Douglas A. Lauffenburger, Linda G. Griffith

## Abstract

Three-dimensional *in vitro* liver models are a promising means to recapitulate key aspects of human liver disease pathologies, thereby aiding therapeutic development. Spheroidal aggregates of hepatocytes, sometimes including non-parenchymal cells, are an established approach for modeling facets of metabolism and drug responses, yet these models often lack dynamic interactions with vascular and immune cells that also contribute to disease development and progression. To address this, we developed a microphysiological system (MPS) that integrates multicellular human hepatic spheroids with self-organized microvascular networks. We show extensive interaction between primary human spheroids and functional vasculature while maintaining key hepatic functions. We demonstrate the utility of this MPS by modeling an insulin resistance state through chronic exposure to disease-mimetic media conditions. This disease model displays altered hepatocyte metabolism, dysregulated vascular features, and increased inflammation state. Enabled by the functional vasculature, we further extend this disease model to capture changes in immune cell recruitment. When culturing CD14+ monocytes in our liver MPS, a subset of monocytes extravasate, localize to hepatic spheroids, and begin differentiating into CD163+ macrophages. These cells infiltrate with greater frequency in insulin resistant samples, consistent with known clinical findings. All together, this vascularized MPS model captures disease-relevant liver biology including inflammatory features.

## Introduction

The liver is responsible for a diverse set of vital functions, spanning plasma protein production, nutrient metabolism, immune regulation, and drug and toxin clearance.^1,2^ This versatility is enabled by both the presence of tissue-specific cell types as well as their complex architectural organization, with hepatocytes acting as the primary arbiter of many of the organ’s key activities. In carbohydrate metabolism, the liver helps to preserve blood sugar homeostasis by maintaining a balance between the storage and secretion of glucose. Additionally, the liver is a central regulator of lipid metabolism, with hepatocytes responsible for breaking down free fatty acids into triglycerides for storage.^1,3^ Given this organ’s essential involvement in mediating metabolic regulatory functions, dysregulation induces systemic changes that culminate in metabolic disorders such as type 2 diabetes (T2D) and metabolic-associated steatotic liver disease (MASLD).^4^

A fundamental element of these diseases is insulin resistance, defined as the body’s inability to respond effectively to the peptide hormone insulin, which plays a key role in regulating energy storage.^5^ In the healthy liver, insulin directly induces glucose uptake and glycogen synthesis, upregulation of lipogenic gene expression, and downregulation of gluconeogenic gene expression. In contrast, insulin resistance results in impaired insulin clearance, aberrant hepatic glucose uptake and production, and elevated lipid synthesis.^6,7^ Importantly, insulin resistance is associated with activation of liver-resident macrophages (Kupffer cells) and elevated pro-inflammatory cytokines such as IL-6 and TNF-α; this immune system response is considered part of a cycle via which inflammation can further drive insulin resistance and recruitment of circulating immune cells, which in turn causes chronic and systemic immune dysregulation.^8,9^

Much remains to be understood about the mechanisms by which insulin signaling in the liver is altered, as well as the complex interactions between hepatic and systemic insulin resistance leading to disease progression. These knowledge gaps are in part due to a historical reliance on animal models, which often exhibit significant disparities from human phenotypes and establish pathophysiology through dissimilar mechanisms.^10,11^ As a result, translation of most findings to the human context has seen limited clinical success. To date, resmetirom is the only U.S. FDA-approved therapeutic for adults with metabolic dysfunction-associated steatohepatitis (MASH), an advanced stage of MASLD, exemplifying the ongoing gap in our understanding of metabolic liver disease.^12^

In recent years, microphysiological systems (MPS) have emerged as a promising alternative to animal models, presenting new opportunities for advancing our understanding of human biology and disease.^13,14^ By enabling 3D co-culture of multiple cell types within meso-or microfluidic devices, MPS can maintain primary human liver phenotypes longer and more faithfully than culture in 2D.^14–17^ Further, efforts to incorporate tissue-specific non-parenchymal cells (NPCs) — including Kupffer cells, stellate cells, and liver sinusoidal endothelial cells (LSECs) — capture key functional contributions and cellular interactions that are otherwise lost in simpler culture contexts.^8,18,19^ Recent studies demonstrating improved primary human hepatocyte longevity and function with either synthetic vascular-like channel structures or co-culture with endothelial cells underscore the value of adding vascular elements.^20–22^ In addition, accessible microfluidic models with 3D perfusable vasculature have been employed to study immune cell trafficking for some applications,^23–26^ but they have not yet been used to enable direct transport of immune populations to hepatocytes themselves in situ, in part due to challenges with integrating perfused vasculature within hepatic tissue structures.^27–30^

In parallel, liver MPS have been applied to a variety of disease states, including MASLD / MASH. These models employ tailored culture media compositions (*e.g*., high nutrient concentrations) designed to induce specific phenotypes (*e.g.,* features of hepatocyte steatosis).^31–35^ While inclusion of NPCs within these disease models has improved their fidelity to disease phenotypes, the known roles of vasculature and immune cell trafficking in disease pathogenesis motivate our building more accessible models that incorporate these elements.^36–39^

Here, we establish a primary human cell-based, vascularized disease model of hepatic insulin resistance. By leveraging and improving upon self-organized *in vitro* vascular models, we developed a microfluidic platform where vasculature perfuses directly into multicellular hepatic spheroids comprising hepatocytes, Kupffer cells, endothelial cells, and fibroblasts, exhibiting high degrees of interactions and enabling transport directly into spheroid structures. Beyond supporting hepatocyte and vascular function, this MPS also captures features of disease phenotypes. A pathological disease media formulation induces an insulin resistant state in this vascularized liver platform, with altered insulin clearance and inflammation profiles. Finally, we demonstrate that these altered inflammation states functionally impact the relationship between the local environment and systemically circulating innate immune cells. With the addition of peripheral monocytes, we show enhanced monocyte recruitment under insulin resistance conditions, recapitulating immune-tissue interactions in metabolic disease.

## Results

### Multicellular hepatic spheroids integrate with self-organized vascular networks

We first generated liver spheroids using an alginate microwell system.^40^ These spheroids comprised donor-matched primary human hepatocytes and Kupffer cells; human umbilical vein endothelial cells (HUVECs); and normal human lung fibroblasts (NHLFs). The formation of perfusable microvascular networks in fibrin hydrogels via morphogenesis of HUVECs and NHLFs is a well-established protocol in the MPS community. We speculated that also including these two vascular elements in our hepatic spheroids would assist in establishing vascular junctions between the spheroids and their surrounding microvasculature.^41,42^ To create a suitable microfluidic platform, we adapted a polydimethylsiloxane (PDMS) device design that is well-established for vascularization studies^43–46^ by bonding devices to 6-well glass bottom plates, facilitating higher throughput culture and improved long-term time-lapse imaging **(SI Fig 1)**. After these spheroids compacted for 2 days, we harvested them, mixed them with fibrin gel precursor containing additional HUVECs and NHLFs, and introduced this mixture into a microfluidic device. After fibrin polymerization, we maintained the cultures with daily media changes for 7-14 days, using a rocker platform to promote vasculogenesis and longer-term maintenance of perfusable microvessels **(Fig 1A)**.

**Figure 1:**
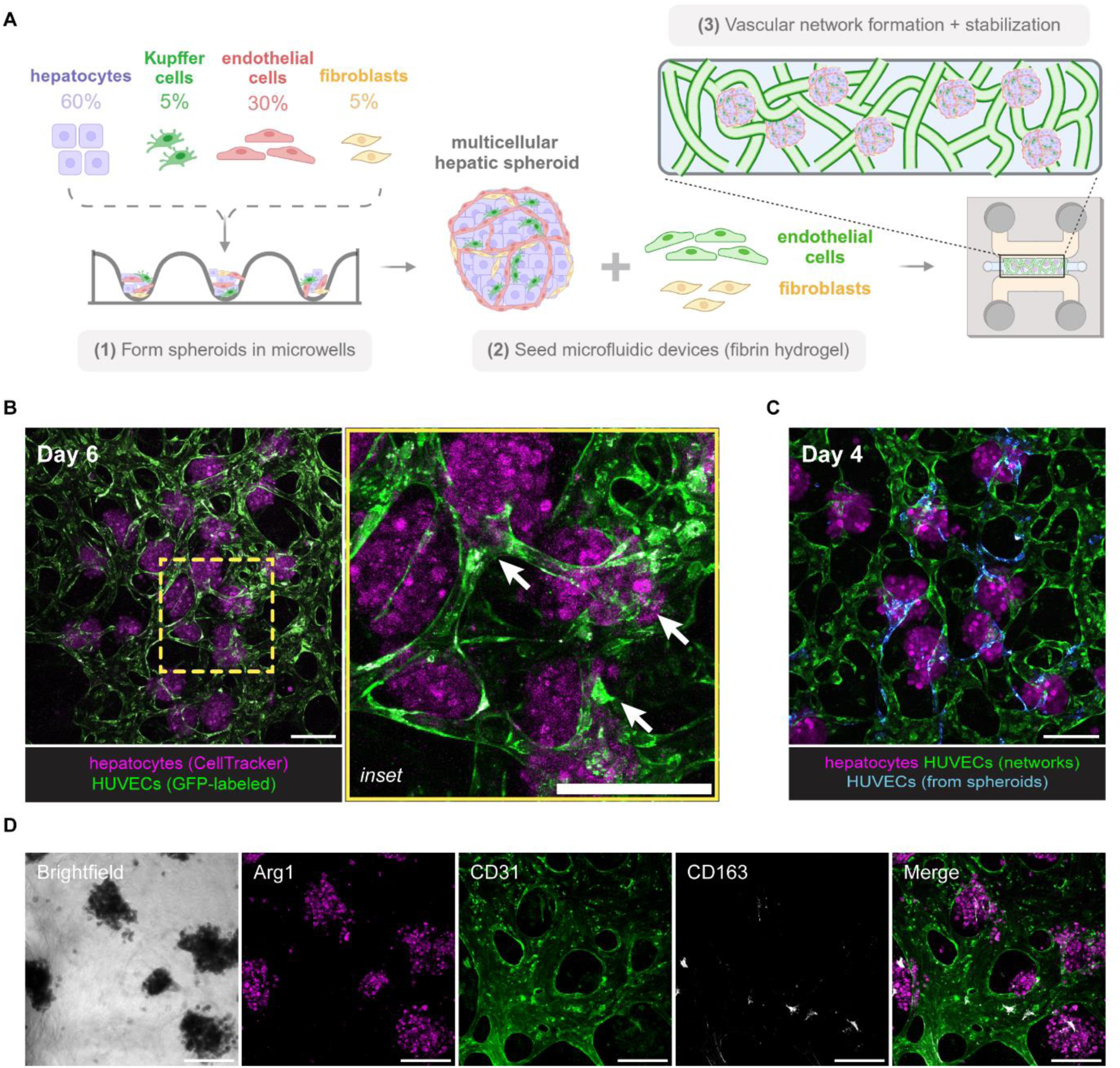
Generation of a vascularized, primary human liver spheroid-based culture platform. (A) Schematic of culture platform and seeding strategy, beginning with multicellular hepatic spheroid generation in alginate microwells for 2 days, followed by harvesting and co-encapsulation of spheroids, endothelial cells, and supporting fibroblasts inside of a fibrin hydrogel within polydimethylsiloxane (PDMS) microfluidic devices. (B) Self-organized vascular network formation after 6 days in devices. Maximum intensity projection of live image taken at 10x magnification shows GFP-HUVEC networks (green) interacting with hepatocytes labeled with CellTracker Deep Red (magenta). Inset (yellow box) shows a magnified view, with arrows indicating regions where networks penetrate through spheroids. Scale bars = 200 µm. (C) Live image taken on Day 4 after device seeding shows RFP-HUVECs (false-colored cyan) migrating out of spheroids to connect with GFP-HUVECs (green), with hepatocytes labeled with CellTracker Deep Red (magenta). Scale bar = 200 µm. (D) Representative maximum intensity projection image of an immunostained device after 14 days of culture. Arginase-1 (magenta) indicates hepatocytes, CD31 (green) indicates HUVECs, and CD163 (white) indicates Kupffer cells. Scale bars = 200 µm.

To enable co-culture of these primary human cell types, we first defined a media supportive of liver and vascular function, inspired by combining commercial formulations for hepatocyte and endothelial cell media, while maintaining physiological ranges of glucose, insulin, and free fatty acid levels^47^ **(SI Table 1)**. Hepatic spheroid structural maintenance and microvascular network formation and perfusability were both independently assessed in this new media formulation over 1-2 weeks in culture **(SI Fig 2)**.

Live imaging demonstrated that HUVECs self-organize into microvascular networks spanning the device tissue compartment over the first 3-5 days in culture **(SI Fig 3, SI Video 1)**. Microvascular networks both surround and penetrate through the hepatic spheroids, establishing the spheroids as both interacting with and integrated within the microvasculature **(Fig 1B)**. To test whether these interactions were in part due to the endothelial cells within the hepatic spheroids forming links with those external to the spheroid, we generated hepatic spheroids containing RFP-HUVECs and monitored their interactions with surrounding GFP-HUVEC vascular networks. We observed that endothelial cells that originated within the spheroids not only connected the spheroids to the immediately adjacent GFP-HUVEC microvessels, but also migrated to distant locations throughout the networks after several days in culture **(Fig 1C, SI Fig 3)**. Additionally, we confirmed that after 14 days of device culture, hepatocytes and Kupffer cells remain integrated with the networks **(Fig 1D)**.

Beyond visualizing these physical interactions in co-culture, we also evaluated critical structural and metabolic functions of the vasculature and hepatocytes. We established that the microvascular networks had open lumens by perfusing fluorescent dextran through each device and observing the dye localization within vessel structures **(Fig 2A)**. Notably, perfusable segments form through the hepatic spheroids **(Fig 2B, SI Video 2)** and span the entirety of the central hydrogel compartment **(SI Fig 4)**. By having vasculature that perfuses directly through hepatic spheroids and thus enables transport into the tissue structures, this system improves upon previous vascular liver models, which often show vasculature adjacent to liver spheroids rather than in an integrated culture format.^27,29^

**Figure 2:**
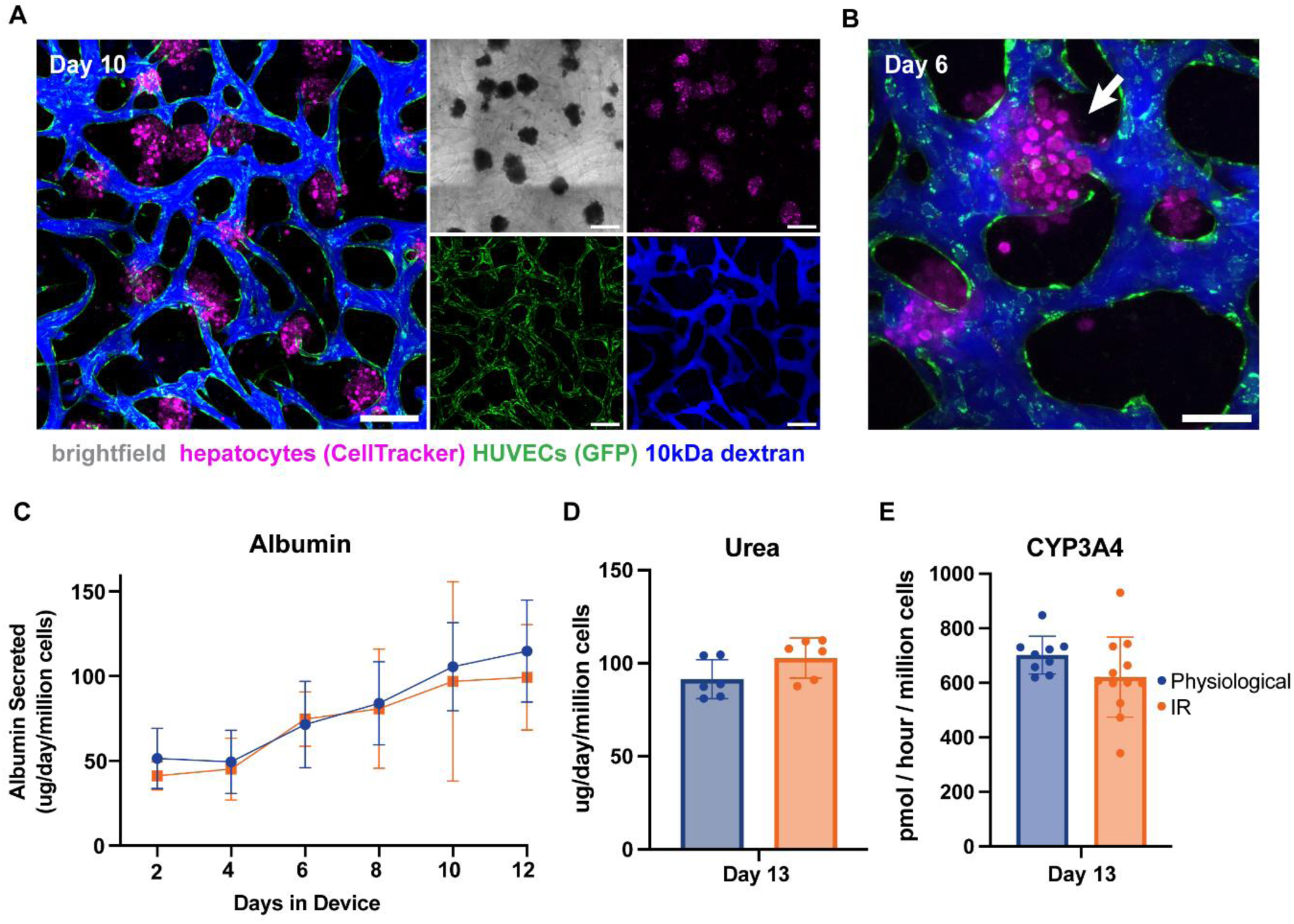
Liver MPS supports the formation of perfusable vascular networks and functional primary human hepatocytes. (A) Maximum intensity projection images of 10kDa dextran (blue) perfusion on Day 10 of device culture demonstrates GFP-HUVEC vascular networks (green) have open lumens. Hepatocytes are labeled with CellTracker Deep Red (magenta). Scale bars = 250 µm. (B) Representative maximum intensity projection image shows that perfusable vessels penetrate through hepatic spheroids on Day 6. Scale bar = 100 µm. (C) Hepatocyte secretion of albumin, normalized per million hepatocytes, shows increased albumin levels over time, n = 11 devices per media condition. (D-E) Hepatocytes maintain urea secretion and Cytochrome P450 3A4 (CYP3A4) metabolism after 13 days in device culture (15 days after initial spheroid formation). Each point represents one device. n = 6-12 devices per media condition.

Hepatocellular function was assessed in both physiological baseline medium and a metabolic overload “insulin resistance”-inducing (IR) media formulation featuring high insulin, glucose, and free fatty acid levels **(SI Table 1)**. Hepatocytes in both media formulations sustained albumin production throughout the course of the experiment, with a drift upward in magnitude over 2 weeks in culture, an improvement over hepatic spheroids alone **(Fig 2C, SI Fig 5)**. Further, hepatocytes exhibit robust CYP3A4 activity and urea production after 2 weeks of culture in MPS devices, with comparable values detected in both media formulations **(Fig 2D-E)**. With these benchmarks of hepatic function meeting ranges in accordance with *in vivo* function^48^, we then investigated additional behaviors of the vascularized liver MPS in simulated disease conditions.

### Vascularized liver platform recapitulates known features of insulin resistance disease state

We specifically sought to understand whether this vascularized liver MPS can be applied to model features of hepatic insulin resistance that contribute to liver and systemic metabolic disorders. We hypothesized that culturing our devices in IR media would induce phenotypes associated with an insulin resistance disease state. We began by investigating the clearance of insulin by the cells in our system, a process known to be dysregulated in hepatic insulin resistance.^7^ While both the physiological and IR media conditions initially showed equal insulin clearance rates, sustained exposure to the IR media resulted in a decline in clearance that is consistent with insulin resistance pathogenesis **(Fig 3A)**.

**Figure 3:**
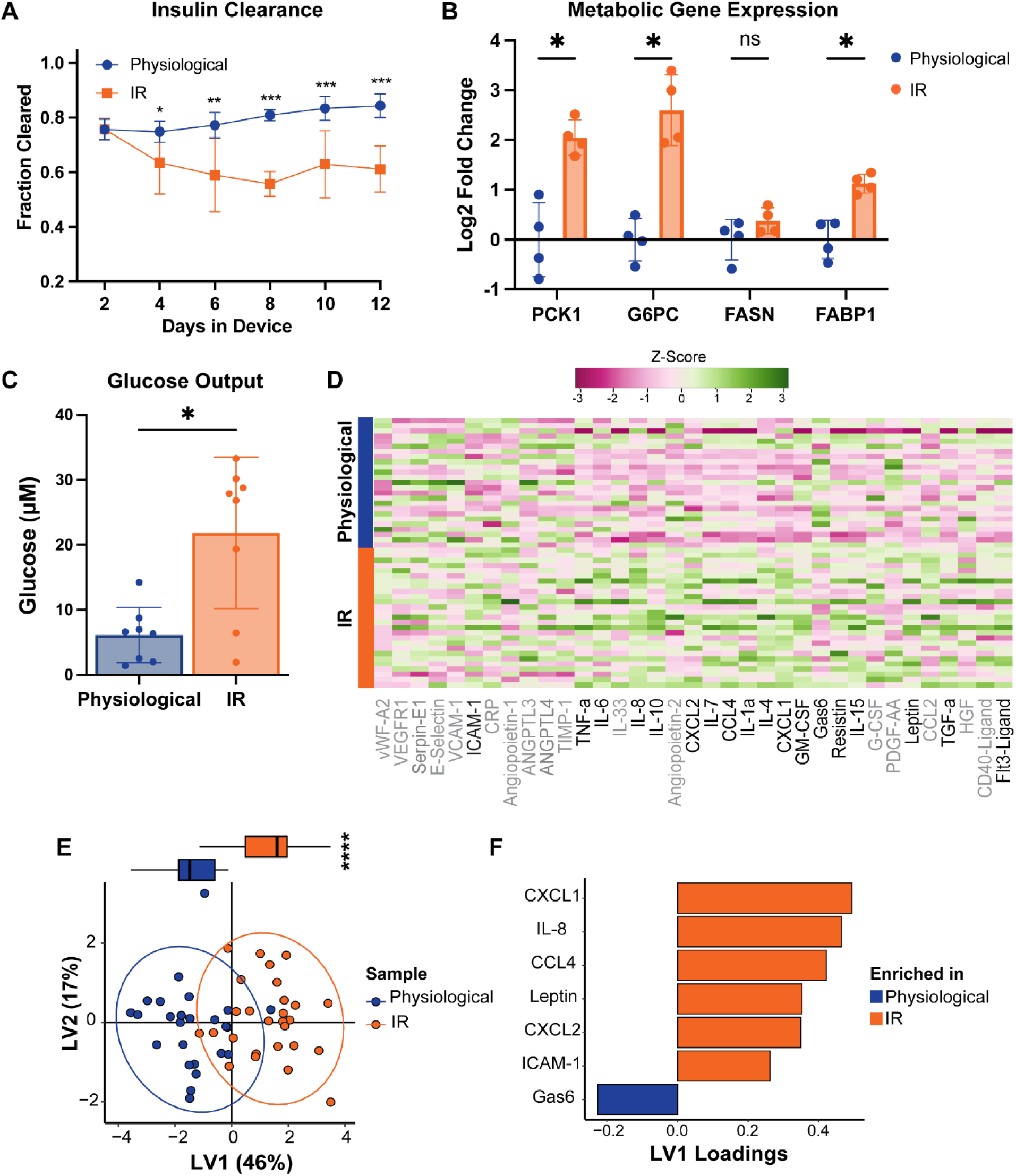
Establishment of an insulin resistance state in vascularized liver MPS. (A) Insulin clearance diminishes over time in devices cultured in the insulin resistance (“IR”) medium formulation, compared to physiological media. n = 8 devices per condition. (B) Gluconeogenesis gene expression (PCK1, G6PC) is significantly increased in IR devices relative to physiological control devices, measured via RT-qPCR. Changes in genes associated with fatty acid transport and synthesis (FASN and FABP1) are upregulated to a lesser degree. (C) Glucose production is increased in IR devices, measured over 24h from day 13 to day 14 of device culture. n = 8 devices per condition. (D) Row-normalized heatmap of cytokines, chemokines, and growth factors assayed via Luminex from cell culture supernatant collected on Day 10 of device culture. Each row represents one device. Analyte labels in black indicate significant differences between physiological and IR conditions (P-adj < 0.05). (E) Partial least squares discriminant analysis (PLS-DA) model built from Luminex dataset separates physiological and IR samples, with each point representing the scores on latent variable 1 (LV1) and LV2 from one device. Ellipses show the 95% confidence interval for each condition, with the box plot quantifying the significance of separation between conditions on LV1. (F) Loadings on LV1 indicate features contributing to the model, with Gas6 associating with physiological conditions, and several chemokines and cytokines associated with the IR condition.

Additionally, insulin signaling partially regulates glucose production by hepatocytes. When comparing the IR condition to the physiological condition, gene expression analysis by qPCR revealed significantly elevated levels of genes involved in gluconeogenesis (PCK1 and G6PC) and smaller changes detected in genes involved in fatty acid transport and synthesis (FASN and FABP1) **(Fig 3B)**. To establish whether this altered gene expression was associated with observable functional changes, we adapted a canonical assay to measure hepatocellular glucose output in a vascularized system, in which media is changed to a reduced glucose production media containing pyruvate and lactate (see Methods) for 24 hours. The IR condition resulted in higher net glucose production compared to the physiological condition, indicating impaired metabolic regulation **(Fig 3C)**. We note that the absolute magnitude of glucose output observed is lower than expected compared to hepatocyte-only 3D culture, likely due to the glycolytic metabolism resulting in glucose consumption by vascular cells in co-culture. Even so, the greater net glucose in IR conditions is consistent with the observed metabolic gene expression changes.

Due to the association between chronic inflammation and insulin resistance, we also analyzed a panel of chemokines, cytokines, and growth factors associated with inflammation state or vascular dysregulation. This panel was assayed via Luminex on supernatant collected on day 10 of device culture **(Fig 3D)**. To better identify key features associated with the IR state, we performed supervised multivariate analysis to identify features associated with each media condition using partial least squares discriminant analysis (PLS-DA). This model was able to significantly separate samples by physiological and IR conditions **(Fig 3E)**. Examining the PLS-DA loadings driving this separation, we see chemokines such as CXCL1, CXCL2, and CCL4 correlating with IR conditions **(Fig 3F)**. This aligns with clinical observations of increased hepatic infiltration by immune cells in IR and MASLD, with this acute inflammation further driving chronic insulin resistance.^49,50^

Interestingly, from our Luminex panel we also saw increased secretion of the glycoprotein ICAM-1 associated with the IR devices, suggesting a pro-inflammatory endothelial cell disease state. Some patient cohort studies have suggested that elevated plasma levels of ICAM-1 and other cell adhesion molecules could be a predictor for people at increased risk of developing T2D and associated cardiovascular complications.^51,52^ To parse the extent to which the vascular networks themselves contribute to the disease phenotypes observed, we also performed experiments separating the two main components of our model, spheroids and microvasculature. Hepatic spheroids alone in physiological and IR conditions confirmed vasculature was not required for disease state development **(SI Fig 5)**. Meanwhile, microfluidic culture of vascular networks alone demonstrated vascular cells contribute to some metrics, such as insulin clearance **(SI Fig 6A)** and inflammation profiles (SI Fig 6C-E), but features such as gluconeogenic gene expression **(SI Fig 6B)** did not change in the absence of liver cells, as expected.

While we initially focused on hepatocyte-specific metrics in our comparisons of the physiological vs. IR media conditions, we also noticed that IR media led to the development of consistently narrower vascular networks in our liver MPS during initial dextran perfusion tests **(Fig 4A)**. This observation, in combination with our PLS-DA findings of a potentially inflamed endothelial state, motivated us to further quantify macro-scale differences in vascular morphology and permeability **(Fig 4B)**. Imaging our vascularized liver MPS, followed by quantification of vessel features^53^, showed that by Day 6, there was significantly reduced vessel coverage under IR conditions compared to the physiological baseline **(Fig 4C)**, as well as narrower vessels **(Fig 4D)**. In parallel, our permeability analysis showed that the narrowed, IR-associated microvascular networks also display increased permeability, resulting in “leakier” vessel structures **(Fig 4E)**. Interestingly, these results not only mimic reported effects of high glucose on 2D endothelial cultures, but also reflect microvascular complications often seen in diabetic patients, with increased capillary permeability and narrower vessels being hallmarks of diabetic microangiopathy and peripheral vascular disease.^54–57^

**Figure 4:**
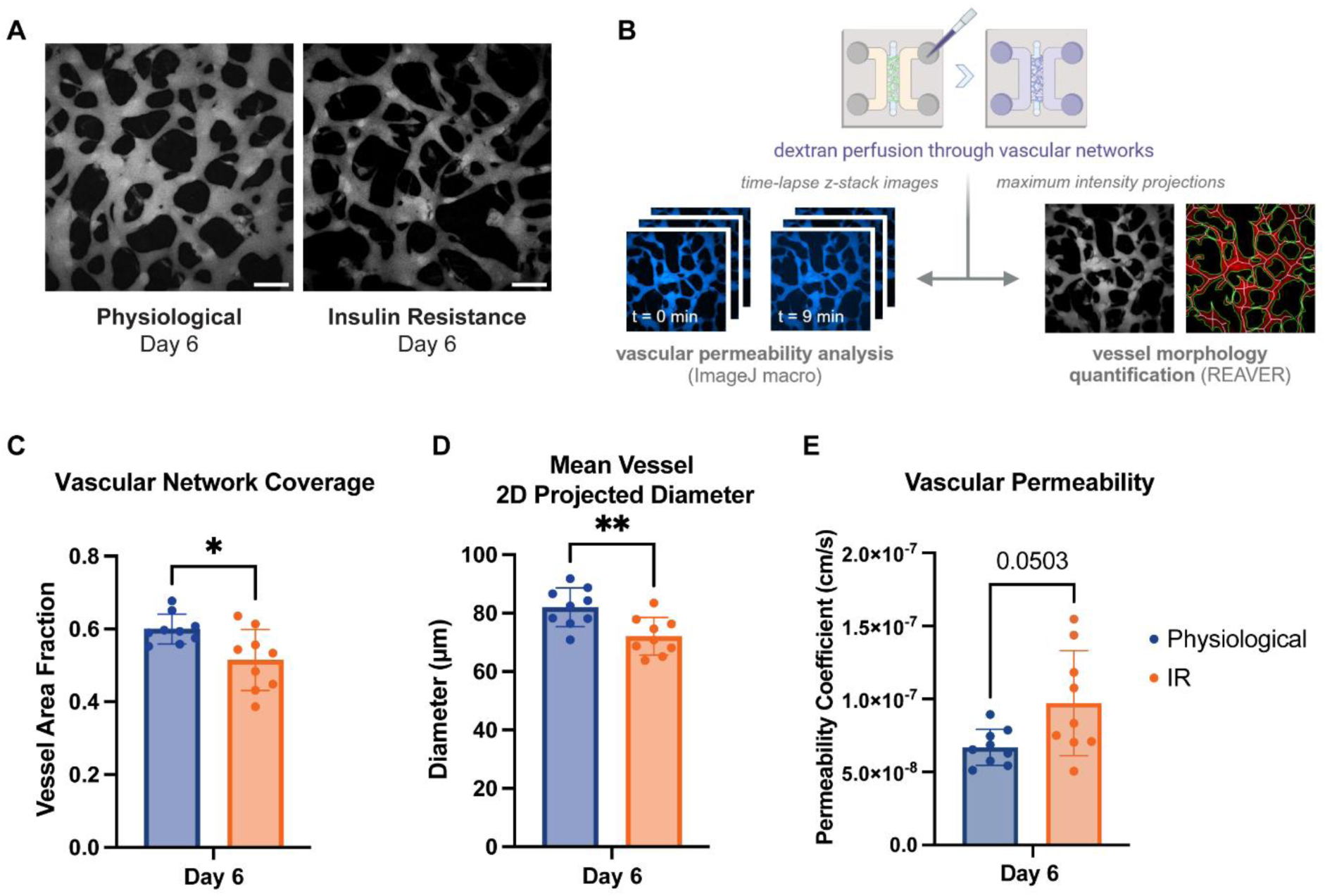
Vascular networks in liver MPS display hallmarks of peripheral disease. (A) Representative images of vascular network morphology after 6 days in culture, with maximum intensity projections of the fluorescent dextran channel shown for visualizing perfusable vessels. Scale bars = 200 µm. (B) Schematic illustrating protocol for imaging vascular networks and performing downstream analyses. Vascular permeability coefficient is calculated using z-stack images taken of the same region of interest at two defined time points, imported into an ImageJ macro. Vessel morphological parameters are quantified using maximum intensity projection images of the same z-stacks, loaded into the REAVER MATLAB tool. (C) Insulin resistance (“IR”) culture condition results in vascular networks that cover a significantly smaller area, compared to the physiological baseline. n = 9 devices per media condition, with quantification from n = 3 ROIs averaged per device. (D) IR culture condition results in vascular networks that have a significantly smaller mean 2D projected diameter (“mean vessel diameter” in REAVER), compared to the physiological baseline. n = 9 devices per media condition, with quantification from n = 3 ROIs averaged per device. (E) Vascular networks in IR devices exhibit increased permeability to 10kDa dextran, with z-stack images (5µm step size, 10 slices) taken 9 minutes apart. n = 9 devices per media condition, with permeability measurements from n = 3 ROIs averaged per device.

### Establishing peripheral immune - liver interactions

In addition to peripheral vascular dysregulation, insulin resistance and MASLD are also clinically associated with chronic inflammation and increased immune cell recruitment to the liver.^49,50^ These features, in combination with our earlier results demonstrating elevated chemokine and cytokine levels in the IR liver MPS, motivated us to investigate immune cell recruitment to hepatic spheroids as a function of induced disease state.

For these experiments, we generated the vascularized liver MPS as previously described in either physiological or IR conditions. After one week, we added CD14+ monocytes freshly isolated from healthy human donors to the microvasculature and maintained this immune co-culture under static conditions for an additional 4 days **(Fig 5A)**. Time-lapse imaging of CellTracker-labeled monocytes showed these cells immediately travel through the vascular networks **(Fig 5B)**. While some quickly flowed to the other side of the devices, others attached to the microvascular networks, extravasated through vessel walls, or localized to hepatic spheroids within the first 12 hours **(Fig 5C)**. After 4 days, we saw these monocytes persist within the vascularized liver MPS, with many remaining closely associated with the hepatic spheroids.

**Figure 5:**
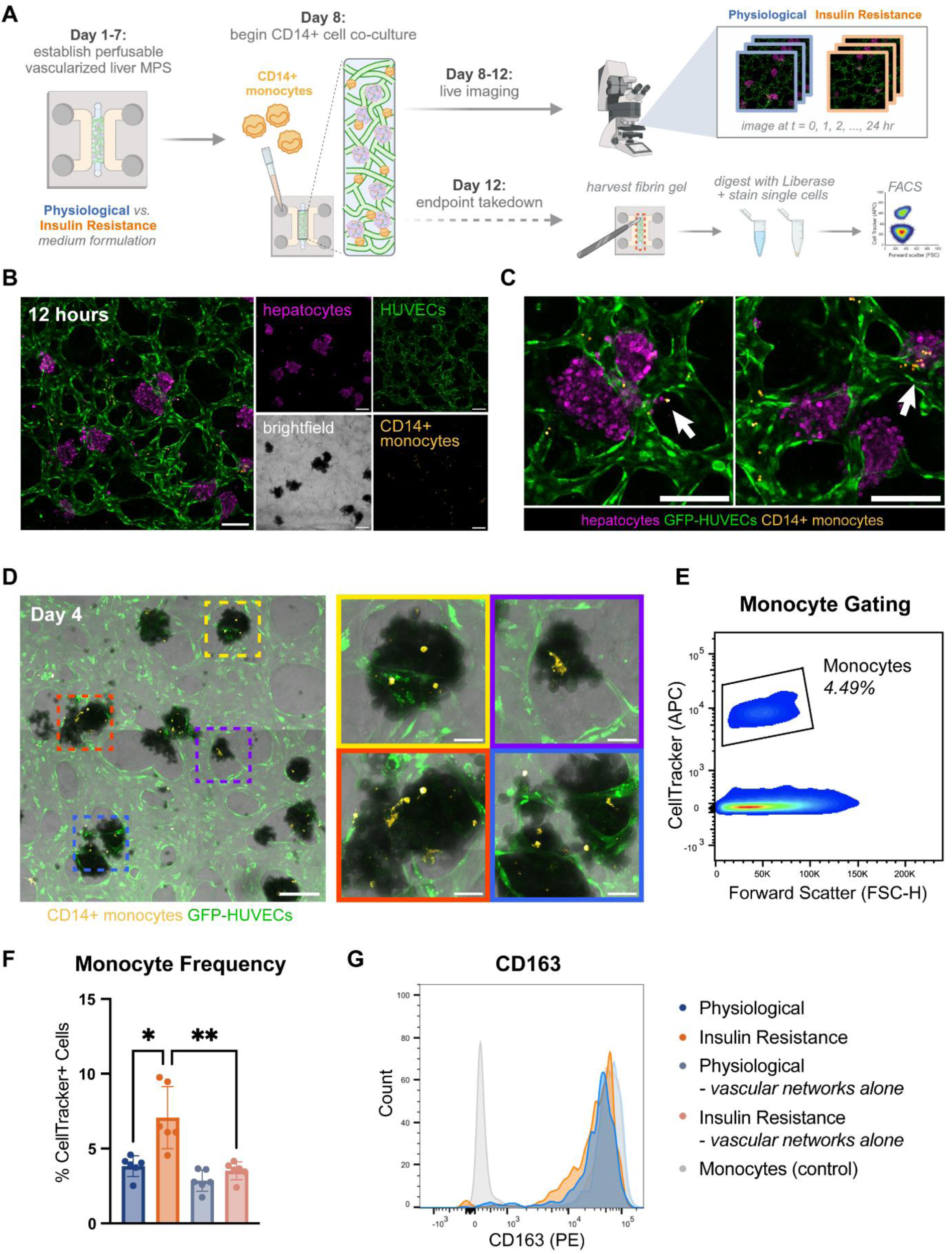
Vasculature supports immune cell trafficking and reveals disease-associated recruitment. (A) Schematic illustrating addition of human PBMC-derived monocytes on day 8 of device culture, after perfusable vasculature has been established, followed by 4 days of co-culture before endpoint fixation or tissue harvesting. (B) Maximum intensity projection of devices 12 hours after monocyte addition shows that monocytes (CellTracker Deep Red, false-colored yellow) localize within GFP-HUVEC vascular networks (green) and begin interacting with spheroids (CellTracker Red, magenta). Scale bars = 200 µm. (C) Representative maximum intensity projection images (insets of 5B) showing monocyte interactions with spheroids. Arrows indicate an example of a monocyte that has extravasated from the vascular networks near a hepatic spheroid (left) and a cluster of monocytes pausing in the networks at a spheroid (right). Scale bars = 200 µm. (D) Imaging after 4 days of monocyte co-culture shows monocytes persist in the devices. Insets (right) show that most monocytes appear to localize with hepatic spheroids, and some monocytes appear to be differentiating based on their changed morphology. Scale bars = 200 µm (left), 50 µm (insets). (E) Flow cytometry gating strategy to quantify the frequency of monocytes (CellTracker Deep Red) in each device. (F) Monocytes were significantly enriched within insulin resistance (IR) devices compared to the physiological condition. Comparison to devices containing vascular networks alone (no hepatic spheroids) also demonstrates the presence of spheroids significantly increases monocyte frequency. n = 6 devices per condition across n = 3 primary monocyte donors. (G) Representative histogram shows elevated CD163 on CellTracker+ monocytes cultured in MPS devices compared to naive monocytes (gray).

Furthermore, a subset of these monocytes appeared to begin differentiating based on their reduced circularity and cell spreading, indicating longer term colonization of the hepatic niche **(Fig 5D)**. To discern if there were significant differences under IR conditions, we performed flow cytometry on these samples to quantify monocyte frequency. To do so, we manually cut out the central compartment from each device, digested the hydrogel to isolate single cells, and ran flow cytometry to measure the fraction of CD14+ cells residing within the tissue compartment of each device **(Fig 5A)**. Gating for the live, CellTracker-positive population in each device allowed for identifying changes in monocyte frequency **(SI Fig 7, Fig 5E)**. We saw significantly increased monocyte frequency in the IR liver MPS samples. To further understand the role of hepatic spheroids in driving this immune cell recruitment, we also cultured monocytes in devices that contained only vascular networks. Comparisons between our full liver MPS model and this vascular networks-only control revealed the hepatic spheroids were responsible for significantly elevated monocyte recruitment **(Fig 5F)**. Cell surface marker characterization revealed monocytes cultured in these devices display elevated CD163 compared to the initial monocytes, indicating differentiation towards CD163+ tissue macrophages **(Fig 5G)**. Previous studies have examined soluble CD163 as a marker of insulin resistance.^58,59^ Future work characterizing the effects of the recruited CD163+ macrophages in our system, including activation and shedding of CD163, could be an interesting avenue for future investigation. Ultimately, these altered frequencies between conditions and monocyte differentiation highlight the ability to examine altered recruitment and immune-tissue interactions in this MPS.

## Discussion

We have established here an MPS model featuring functional primary human liver cell aggregates co-cultured with a perfusable microvasculature surrounding and permeating the hepatic spheroids. We have further demonstrated that this model can be driven by culture media modifications to recapitulate certain metabolic and immune behaviors observed in the insulin-resistant state: metabolic dysregulation, inflammation, and macroscale vascular dysfunction. This modeling was enabled by employing a physiologically relevant media formulation with insulin, glucose, and free fatty acid concentrations in line with patient values. Notably, by starting with this clinically informed baseline we enhance our ability to interrogate subsequent disease state perturbations that more closely mimic human biology — especially in comparison to traditional culture models, which often contain supraphysiological concentrations of nutrients and insulin that far exceed even pathological levels.

We note that this model of insulin resistance most closely represents an earlier phase of disease, without signs of cell death and lipotoxicity for example, which may be observed as insulin resistance progresses to MASLD / MASH. With its modular media components and cell ratios, however, this liver MPS could also be adapted to model more advanced disease, where different parameters could be tuned to mimic and investigate a wider spectrum of MASLD. For example, using primary cells from diseased donors or further altering nutrient, hormone, and/or cytokine levels could recapitulate later metabolic disease stages featuring fibrosis and severe steatosis, though such alterations may also require maintaining cultures for much longer than the two weeks described here.

In addition, this MPS could be applied to study liver-specific vascular features that accompany and contribute to metabolic dysregulation^60^, using LSECs and stellate cells in the spheroids instead of HUVECs and NHLFs, and liver-derived large vessel endothelial cells for the external vasculature. Such studies of vascular function would also require continuous, unidirectional flow and replacement of fibrin with a synthetic extracellular matrix that more robustly withstands cell-mediated remodeling and flow conditions — modifications we are currently implementing.

Our studies of monocyte recruitment were facilitated by the highly perfusable vascular networks that fully integrated with our hepatic spheroids. While it has been clinically established that there is increased monocyte recruitment in metabolic liver diseases such as MASLD and MASH, the timing and relationship between the recruitment of this immune population and a loss of tissue-resident Kupffer cells remains incompletely understood.^39,61,62^ Our model is poised for investigating such phenomena, and others, through further studying downstream changes in recruited immune cell states.

Finally, it is important to highlight that when studying multicellular models, bulk measurements represent a net response of the full system that is influenced by each cell type. For example, in our liver MPS we saw that the vascular networks themselves not only exhibit disease states, but also contribute to broader phenotypes. While incorporating a greater number of cell types may ultimately improve physiological relevance of the system, it also complicates interpretations of experimental results, requiring additional specialized assays to parse cell type-specific contributions, especially given the inherent variability across devices. One approach to further examine this interplay between tissue and vasculature *in vitro* is through altering the ratio of hepatic spheroids to vascular network cells in our model. Initial experiments varying this ratio indicated it was an accessible approach to better infer spheroid contributions, though more extensive optimization will be required to define an optimal spheroid ratio without introducing undue vascular heterogeneity **(SI Fig 8)**.

In conclusion, we present a liver MPS model of perfused, vascularized primary human hepatic spheroids. This platform recapitulates key features of hepatic insulin resistance, with demonstrated infiltration of monocytes further establishing a system capable of modeling complex liver disease biology and potentially guiding future therapeutic development.

## Methods

### Cell Culture

Human umbilical vein endothelial cells (HUVECs; Angioproteomie, cAP-0001) and normal human lung fibroblasts (NHLFs; Lonza, CC-2512) were cultured in flasks using VascuLife VEGF Endothelial Medium Complete Kit (VascuLife; LifeLine, LL-0003) and FibroLife S2 Fibroblast Medium Complete Kit (FibroLife; Lifeline, LL-0011), respectively. GFP- and RFP-expressing HUVECs were used in studies that involved real-time visualization of microvascular network formation (Angioproteomie, cAP-0001GFP and cAP-0001RFP). Cells were cultured at 37°C and 5% CO2 in a humidified incubator, with media changed every other day. HUVECs and NHLFs were detached using TrypLE (ThermoFisher) and used between passages 5-9 for all experiments.

Primary human hepatocytes and Kupffer cells in this study were donor-matched from a 53-year-old Hispanic male (LifeNet Health LifeSciences). Hepatocytes were thawed in Cryopreserved Hepatocyte Recovery Media (CHRM, ThermoFisher), spun down at 100g for 8 minutes, and resuspended in our custom “physiological” William’s E / Vasculife media formulation **(SI Table 1)**. Kupffer cells were thawed in ice cold William’s E media and spun at 450g for 5 minutes before being resuspended in physiological medium. Both hepatocytes and Kupffer cells were thawed directly before spheroid formation.

### Multicellular Hepatic Spheroid Formation

Spheroids were formed using alginate microwells as previously published.^40^ Alginate microwells were primed with 5% BSA for at least 2 hours at 37°C before seeding. 120,000 primary human hepatocytes, 60,000 HUVECs, 12,000 Kupffer cells, and 12,000 NHLFs were pre-mixed to a volume of 75µL per well. After removing the BSA solution, alginate microwells were primed with 75µL of culture media, then the spheroid cell mixture was added dropwise directly onto the alginate microwells. Wells were spun at 50g for 2 minutes before being topped with 150µL additional media. Media was refreshed after 24 hours. For studies that involved live imaging, hepatocytes were labeled with CellTracker Deep Red (ThermoFisher, C34565) after thawing and before addition to alginate microwells, at 1mM concentration according to manufacturer instructions.

### Microfluidic Device Fabrication

The microfluidic devices used in this study consist of three parallel microchannels and were fabricated using soft lithography as previously described.^44^ Briefly, polydimethylsiloxane (PDMS; Sylgard 184, Ellsworth Adhesives) elastomer and curing agent were mixed in a 10:1 w/w ratio, degassed and poured onto a silicon master mold, and cured at 60°C overnight. The PDMS was removed from the mold and cut into individual devices, and biopsy punches were used to create inlet/outlet ports in the central gel channel (2mm) and media reservoirs (5mm). Dust was removed from the PDMS surfaces using Scotch tape before the devices were dry sterilized in the autoclave. The PDMS devices were then treated with plasma (Harrick Plasma Cleaner) for 2 minutes and bonded to 6-well glass-bottom plates (Cellvis, P06-1.5H-N) to facilitate efficient live imaging during experiments **(SI Fig. 1)**. After plasma bonding, devices were left overnight in a 60°C oven to recover hydrophobicity before use. For this study, a device geometry was selected in which the central gel channel measures 3mm wide and 500µm tall, in order to provide sufficient space for microvascular networks to form around the ∼150µm spheroids.^43^ For experiments aimed at testing a higher ratio of spheroids to vascular network cells, a smaller microfluidic device geometry was used, with a central gel channel that measures 1.8mm wide and 400µm tall and accommodates a total of 10µL of gel. For initial media screens to test perfusable network formation (without spheroids), AIM Biotech idenTx chips were used with the same cell concentrations.

### Device Seeding

Spheroids were harvested from alginate wells by pipetting with a wide bore pipette tip and incubated with AlgiMatrix Dissolving Buffer (Gibco, A11340) for 5 minutes before spinning down at 50g for 2 minutes. Supernatant was aspirated and cells were resuspended in 1mL of media. After spheroids naturally settled by gravity, media was aspirated to remove loose single cells and spheroids were resuspended again before being combined with HUVECs and NHLFs.

Fibrinogen from bovine plasma (Sigma, F8630) was dissolved at 37°C in Dulbecco’s Phosphate-Buffered Saline (DPBS) to a concentration of 6 mg/mL. Thrombin from bovine plasma (Sigma, T4648) was reconstituted to 100 U/mL in 0.1% w/v bovine serum albumin in water, and then diluted in physiological media to a concentration of 3- 4 U/mL. HUVECs, NHLFs, and spheroids were combined in a master mix and centrifuged at 200g for 5 minutes, with the resulting cell pellet resuspended in the diluted thrombin solution. For each device, 15µL of the thrombin cell suspension was mixed thoroughly with 15µL fibrinogen (final concentration: 3 mg/mL) and pipetted into the central gel channel using a wide bore pipette tip. The final concentrations of HUVECs, NHLFs, and spheroids were 7M/mL, 1.5M/mL, and ∼200 spheroids/device, respectively. Some devices included only HUVECs and NHLFs as a “vascular networks alone” control. After seeding, devices were moved to a humidified incubator for 15 minutes to allow the fibrin gel to polymerize, after which cell culture medium was added to the side channels and reservoirs. To provide interstitial flow to aid with vasculogenesis, devices were placed on an OrganoFlow rocker platform (Mimetas) set to tilt 8° every 8 minutes along the axis perpendicular to the gel channel during the experiment.

### Device Culture

Media changes (150µL removed from and added to the reservoirs) were performed every 24 hours. Collected media was cleared of cell debris by spinning at 1000g for 5 minutes immediately after collection, then the supernatant was frozen and stored at - 80°C until further use. Four days after initial device seeding, a monolayer of HUVECs was added (1 million/mL, 30µL per side channel) as described previously, in order to facilitate the generation of perfusable microvascular networks and to prevent non-luminal transport of dye or immune cells between the central gel compartment and media channels in downstream assays.^44^

### Albumin Quantification

To assess hepatocyte spheroid function, stored cell supernatant was thawed at 4°C and albumin secretion was assayed via ELISA (Bethyl Laboratories, E80-129) according to manufacturer instructions.

### CYP3A4 Assay

Devices were rinsed with PBS before adding media containing Luciferin-IPA reagent at a 1:1000 dilution from CYP3A4 P450-Glo™ kit (Promega). Devices were incubated for 1 hour before media collection and storage at -80°C until downstream analysis. Media was thawed at room temperature and the assay was performed according to manufacturer protocol alongside a luciferin standard curve to calculate CYP3A4 levels. Plates were read with a Spectramax i3x plate reader luminescence cartridge.

### Insulin Clearance Quantification

Insulin clearance was evaluated from cell supernatant via ELISA (R&D, DY8056). Clearance fraction was calculated by dividing the measured insulin concentration in cultured media by the measured insulin concentration in cell-free control wells.

### Glucose Output Assay

To assess glucose production, devices were rinsed 3 times for 10 minutes in William’s E media containing no serum, glucose, or insulin. Next, devices were transitioned to a “Glucose production media” containing 50µm glucose, 20mM lactate, and 1mM pyruvate for 24 hours. (William’s E basal medium, 3.5% dialyzed Heat Inactivated FBS, 2.5ng/mL VEGF, 2.5 ng/mL EGF, 2.5ng/mL FGF-b, 7.5 ng/mL IGF-1, 2mM Glutamax, 1% Pen/Strep, 100nM hydrocortisone, 7.5mM HEPES, 50µm glucose, 20mM lactate, and 1mM pyruvate) Collected media was spun down at 1000g for 5 minutes. Glucose concentration was then measured with the Amplex™ Red Glucose Assay kit (A22189, Invitrogen).

### Luminex Assay

Supernatant was analyzed according to manufacturer instructions using R&D Systems Custom 28-plex or 12-plex analyte panels. Assay kits were adapted to fit a 384 well format, using 12.5uL of sample and antibody beads per well. Samples were run at both low (2X) and high (25-30X) dilutions in duplicate on a Bio-Plex 3D suspension array system (Bio-Rad). Downstream analysis was performed in MATLAB to fit a 5-parameter logistic standard curve. Any samples with bead count less than 30 or analytes outside the limits of detection were excluded from downstream analysis. Heatmaps were generated on row z-scores using gplots in R (4.0.3). Significance was assessed via Wilcoxon rank-sum test with Benjamini-Hochberg p-value adjustment for multiple hypothesis correction. Partial least squares discriminant analysis (PLS-DA) was performed using ropls (1.20.0) with LASSO feature selection.

### Gel Harvesting

The devices were rinsed 1X with PBS, the PDMS was cut along the media channels with a sterile scalpel, the PDMS above the central gel channel was lifted, and then the gel was scooped into a tube containing 50µL of Liberase^TM^ at 0.5mg/mL in William’s E medium. Gels were then incubated for 20 minutes at 37°C with intermittent mixing by gently pipetting up and down until the gel flowed easily and visibly broke apart. The samples were then spun down at 400g for 5 minutes. Supernatant was removed and cells were transferred to a 96-well plate for staining or frozen in appropriate reagent for downstream RNA or protein extraction.

### RNA Analysis

Harvested cells were frozen in Trizol at -80°C until ready for RNA isolation. Samples were thawed and incubated at room temperature for 5 minutes, then spun down at 1000g for 5 minutes to remove any residual debris. RNA was isolated with Direct-zol RNA MiniPrep kit (Zymo Research) according to manufacturer instructions, including the in-column DNase I treatment step. RNA was quantified using Nanodrop One and converted to cDNA using the High-Capacity RNA-to-cDNA Kit (ThermoFisher Scientific, 4387406).

### Immune Cell Perfusion

Buffy coats were acquired from Massachusetts General Hospital. The blood was diluted 1X in PBS and layered over Lymphoprep (StemCell Technologies), then spun for 30 minutes with no brake. The PBMC layer was carefully removed, spun for 10 minutes, and resuspended in FACS buffer (PBS, 2% Heat Inactivated FBS, 2mM EDTA) before proceeding directly to CD14+ cell isolation using positive selection beads (StemCell, 17858). CD14+ cells were labeled with CellTracker Deep Red (ThermoFisher, C34565) or CellTracker Red CMTPX (ThermoFisher, C34552) for 20 minutes. After resuspending monocytes to a concentration of 1 million/mL, cells were perfused through microfluidic devices by removing all media from the device, followed by addition of 40,000 monocytes to one media channel. Media was allowed to equilibrate, enabling flow of monocytes across the gel channel to the opposing media channel for 5 minutes, before adding an additional 100µL of media to each media channel in static culture, with daily media changes for 2-4 more days.

### Flow Cytometry

Immediately after being harvested from devices, cells were rinsed 1X with PBS and spun at 200g for 5 minutes. Cells were labeled with a fixable viability dye (BioLegend, 423107; BioLegend 423105) for 15 minutes at room temperature. After washing in FACS buffer, cells were incubated with an FC block for 15 minutes before being stained for 40 minutes on ice. Cells were fixed with 4% paraformaldehyde for 15 minutes and stored in FACS buffer at 4°C until measurement on a BD FACSymphony A3 Cell Analyzer (BD Biosciences).

### Vascular Permeability Analysis

After the microvascular networks became perfusable, vascular permeability was measured using time-lapse confocal imaging (Zeiss LSM 880) as previously published.^44^ Briefly, all media was removed from the devices and replaced with a solution of 0.1 mg/mL 10kDa Cascade Blue dextran (ThermoFisher, D1976) in media. Three random, non-overlapping regions of interest (ROIs) were chosen for each device; for each ROI, two z-stacks (step size = 5µm, 10 steps, resolution = 640 x 640 pixels, ROI dimensions >= 600µm x 600µm) were captured 9 minutes apart. Vascular permeability values were computed after a custom FIJI macro was used to calculate morphological parameters and changes in the average fluorescence intensity in the vasculature and matrix over time, as previously described.

### Vessel Morphology Analysis

Maximum intensity projection images were generated from the same ROIs of dextran-perfused devices used in the vascular permeability analysis. Quantitative analyses of microvascular network morphology were performed for these ROIs using the open-source tool REAVER to obtain metrics such as “mean segment diameter” (2D projected diameter) and “vessel area fraction”.^53^

### Immunostaining

Devices were washed 1X with DPBS and fixed in 4% paraformaldehyde for 30 minutes at room temperature on a rocker platform. After fixation, devices were washed 3X with DPBS and permeabilized with 0.5% Triton-X-100 in PBS for 2 hours at room temperature. Devices were then washed and incubated with blocking buffer overnight (5% donkey serum in PBS) at 4°C. Samples were then incubated with some combination of primary antibodies for CD31 at 1:100 (AF806, Biotechne), CD68 at 1:200 (14-0688-82, Invitrogen), CD163 at 1:100 (AF1607, Biotechne), and Arginase-1 at 1:200 (14-9779-82, Invitrogen), suspended in blocking buffer and incubated for 48 hours on a rocking platform at 4°C, before being washed 3 times and incubated with secondary antibodies at a 1:200 - 1:500 dilution (AlexaFluor, Thermofisher) in blocking buffer for 48 hours at 4°C. Images were taken on a Zeiss LSM 880 confocal microscope or a Keyence BZ-X800 fluorescence microscope.

### Statistical Analysis

Statistical analysis was performed using GraphPad Prism 10. Bar plots show averages with error bars representing standard deviation (SD). Mann-Whitney tests were performed in comparisons of two groups. For multiple groups, two-way ANOVAs with appropriate post-hoc tests were performed. Statistical significance was defined as *p < 0.05, **p < 0.01, ***p < 0.0001, and ****p < 0.0001.

## Acknowledgments

This work was funded by NovoNordisk via a sponsored research agreement with the Massachusetts Institute of Technology. E.N.T., E.L.K., and A.D. were each supported by the National Science Foundation Graduate Research Fellowship Program (NSF GRFP) under Grant No. 2141064. E.N.T. was also supported by the Siebel Scholars Foundation. The authors would like to thank Dr. Sif Groth Rønn, Dr. Sara Toftegaard Hjuler, Dr. Rachelle Prantil Baun, Dr. Damien Demozay, and Dr. Dominik Reinhard Pfister from NovoNordisk for constructive discussions. The authors would also like to thank Dr. Jacob Jeppesen, Dr. Shun Zhang, Marie Floryan, Dr. Zhengpeng Wan, Dr. Sarah Shelton, Dr. Lauren Pruett, Dr. Laura Bahlmann, Dr. Priyatanu Roy, and Dr. Brian Joughin for their technical expertise and support. Finally, the authors thank Dr. Jose Cadavid for his feedback on the manuscript.

## Author Contributions

E.N.T., E.L.K., A.J.W., and L.G.G. conceived the study. L.G.G., D.A.L., and R.D.K. supervised the study. E.N.T. and E.L.K. designed, set up, and maintained experiments. K.K.M. and E.L.K. prepared microfluidic devices. E.L.K. and E.N.T. performed imaging. E.N.T. and K.K.M. performed hepatic function measurements. E.N.T. performed and analyzed liver disease assays. E.L.K. and K.K.M. performed and analyzed vascular assays. E.N.T. and A.D. performed immune perfusion experiments. E.N.T. and E.L.K. wrote the manuscript. All authors reviewed and provided feedback on the manuscript.

