## Supplemental Material for "A Vascularized Liver Microphysiological System Captures Key Features of Hepatic Insulin Resistance and Monocyte Infiltration"

Table 1.1
Physiological Media

| Component | Amount | Unit |
|---|---|---|
| William's E (custom - no glucose, no phenol red) | 50 | % |
| VascuLife Basal Medium | 50 | % |
| VEGF | 2.5 | ng/mL |
| EGF | 2.5 | ng/mL |
| FGF-basic | 2.5 | ng/mL |
| IGF-1 | 7.5 | ng/mL |
| Ascorbic acid | 25 | ug/mL |
| Hydrocortisone hemisuccinate | 540 | nM |
| GlutaMAX | 2 | mM |
| Heparin sulfate | 0.09375 | U/mL |
| FBS | 3.5 | % |
| Gentamicin | 15 | ug/mL |
| Amphotericin B | 7.5 | ng/mL |
| HEPES | 7.5 | mM |
| Glucose | 5.5 | mM |
| Insulin | 200 | pM |
| PenStrep | 0.5 | % |
| Linoleic Acid | 20 | uM |

Table 1.2
Insulin Resistance Media

| Component | Amount | Unit |
|---|---|---|
| William's E (custom - no glucose, no phenol red) | 50 | % |
| VascuLife Basal Medium | 50 | % |
| VEGF | 2.5 | ng/mL |
| EGF | 2.5 | ng/mL |
| FGF-basic | 2.5 | ng/mL |
| IGF-1 | 7.5 | ng/mL |
| Ascorbic acid | 25 | ug/mL |
| Hydrocortisone hemisuccinate | 540 | nM |
| GlutaMAX | 2 | mM |
| Heparin sulfate | 0.09375 | U/mL |
| FBS | 3.5 | % |
| Gentamicin | 15 | ug/mL |
| Amphotericin B | 7.5 | ng/mL |
| HEPES | 7.5 | mM |
| **Glucose** | **11** | **mM** |
| **Insulin** | **800** | **pM** |
| PenStrep | 0.5 | % |
| **Linoleic Acid** | **25** | **uM** |
| **Oleic Acid** | **45** | **uM** |
| **Palmitic Acid** | **30** | **uM** |

**SI Table 1: Physiological and insulin resistance media formulations for supporting vascularized liver MPS culture.**

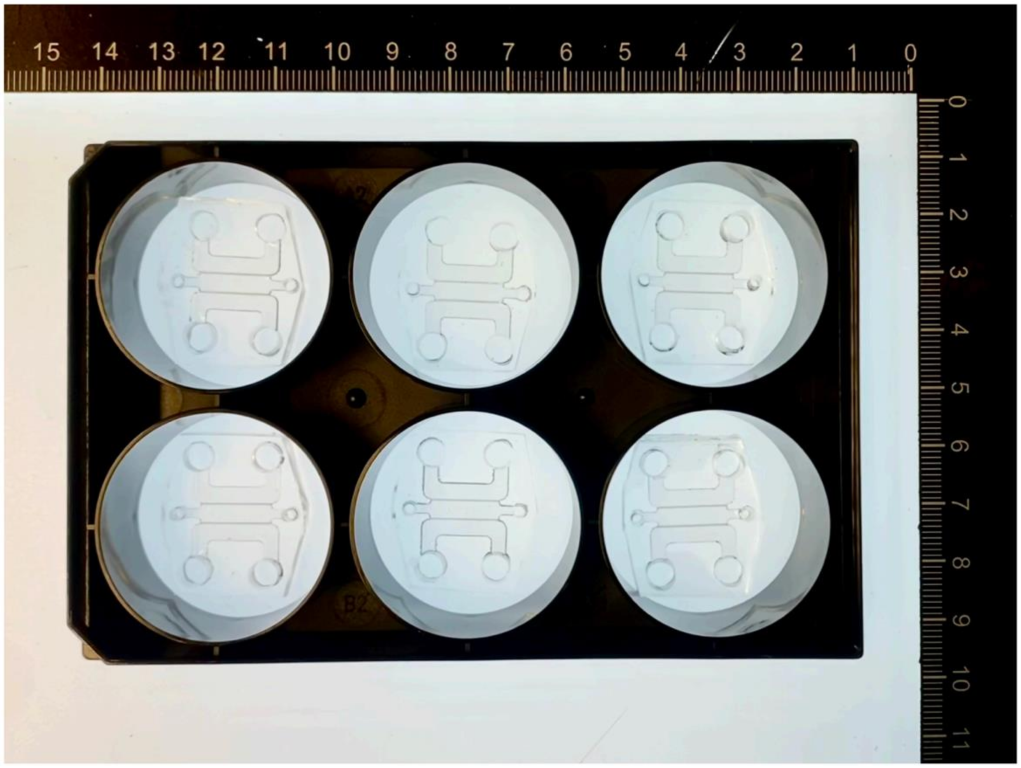


**SI Figure 1: PDMS microfluidic devices are bonded to 6-well glass-bottom plates.** After fabrication, polydimethylsiloxane (PDMS) devices are plasma-treated and then bonded to 6-well glass-bottom plates. Device culture in this plate format allows for increased throughput and accuracy of imaging-based assays, compared to using devices bonded to loose glass coverslips or slides.

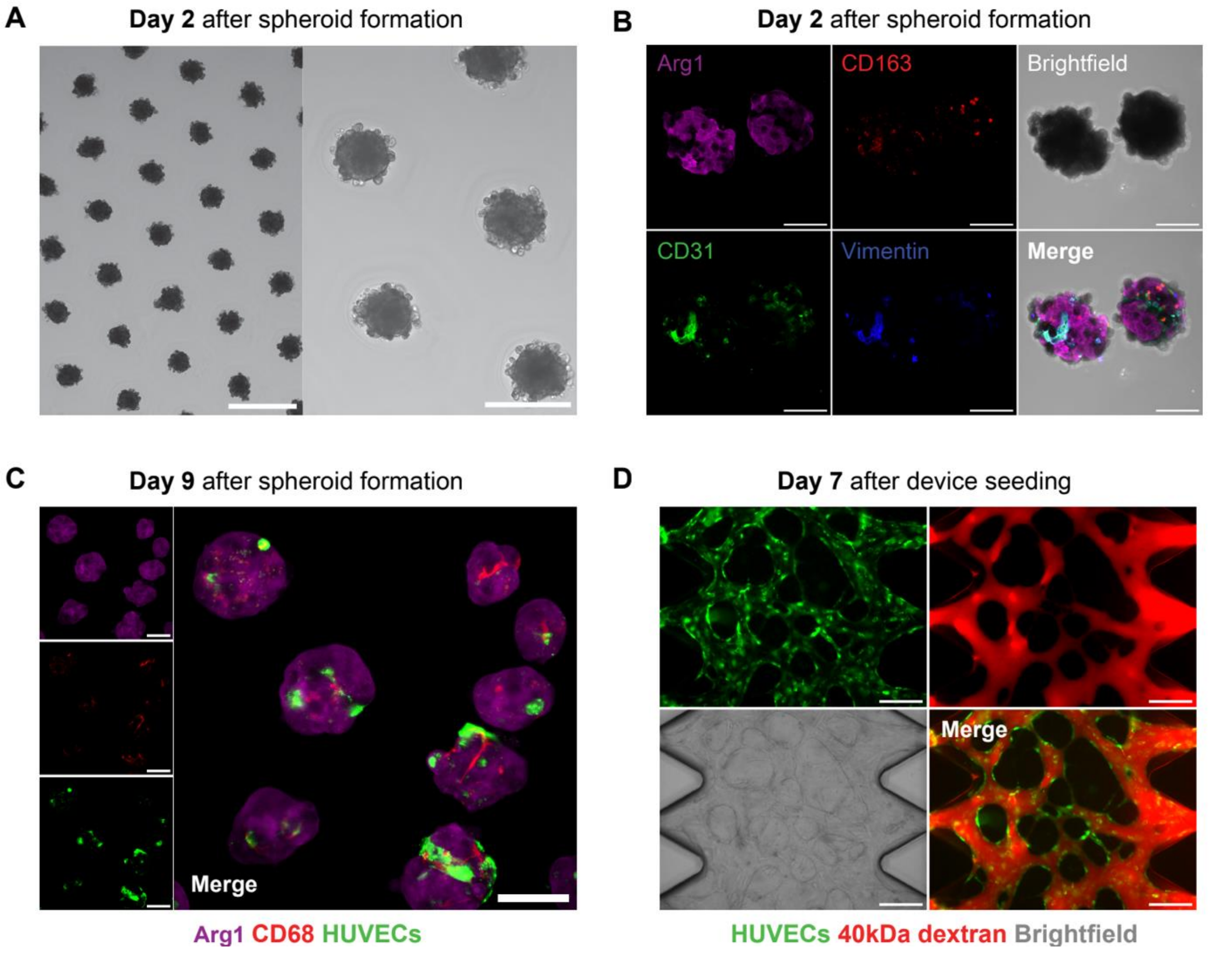


**SI Figure 2: Custom "physiological" media supports formation of multicellular hepatic spheroids and perfusable vasculature.**

A: Spheroid compaction on day 2 in alginate microwells seeded with primary human hepatocytes, primary human Kupffer cells, human umbilical vein endothelial cells (HUVECs), and normal human lung fibroblasts (NHLFs). Scale bars = 500 µm (left) and 250 µm (right).

B: Immunostaining of spheroids 2 days after formation, with all cell types labeled: hepatocytes (arginase-1, magenta), Kupffer cells (CD163, red), HUVECs (CD31, green), and NHLFs (vimentin, blue). Scale bars = 100 µm.

C: Visualization of spheroids 9 days after formation. Hepatocytes are indicated by arginase-1 (magenta), Kupffer cells are indicated by CD68 (red), and HUVECs are RFP-tagged (false-colored green). Scale bars = 100 µm.

D: Networks formed from GFP-HUVECs (green) and NHLFs (unlabeled) are perfusable after 7 days of culture in physiological media in an AIM Biotech idenTx chip, as shown by addition of 40kDa dextran (red) to one media channel. Scale bars = 250 µm.

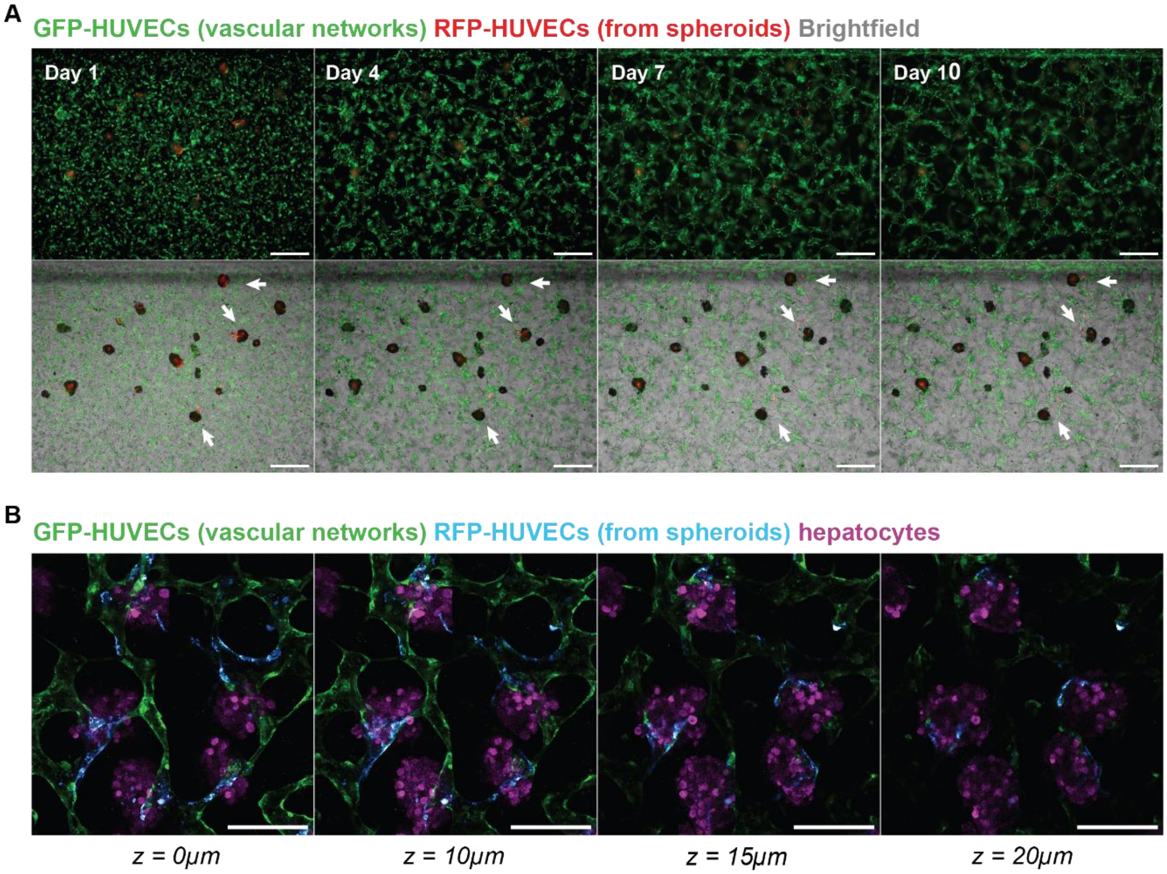


**SI Figure 3: Vascular networks self-assemble around and interact with hepatic spheroids.**

A: Widefield fluorescence images taken on Days 1, 4, 7, and 10 after device seeding show that GFP-HUVECs (green) self-organize into vascular networks over time. White arrows indicate regions where GFP-HUVEC networks are connecting with RFP-HUVECs (red) that migrate out from hepatic spheroids. Overlay of the brightfield channel (bottom row) shows spheroids. Scale bars = 250 µm.

B: Confocal z-slice images (spanning 20 µm in total height) of a Day 4 device show RFP-HUVECs (false-colored cyan) migrating out of spheroids to connect with GFP-HUVECs (green), with hepatocytes labeled with CellTracker Deep Red (magenta). Scale bar = 200 µm.

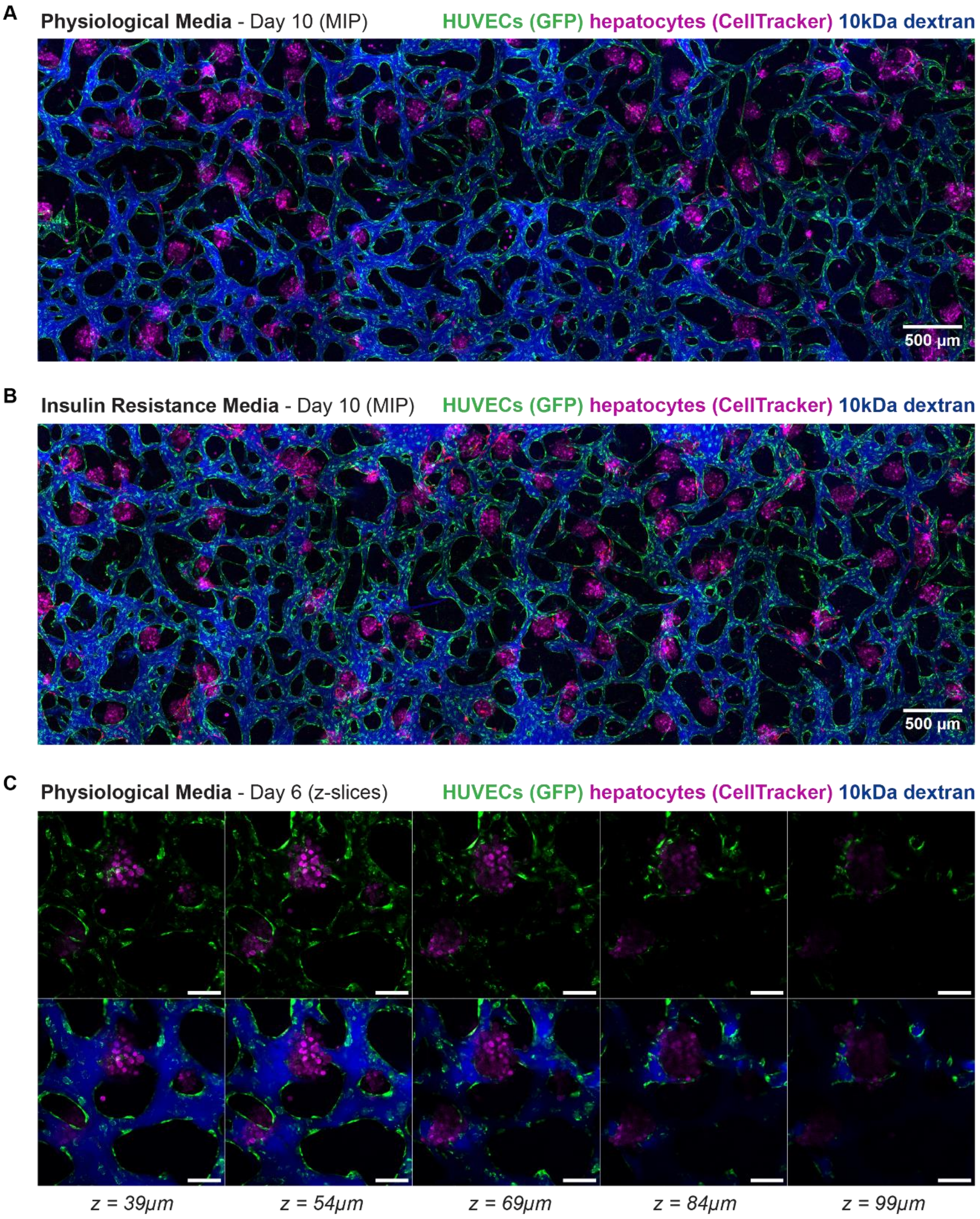
A
Physiological Media - Day 10 (MIP)
HUVECs (GFP) hepatocytes (CellTracker) 10kDa dextran
500 µm
B
Insulin Resistance Media - Day 10 (MIP)
HUVECs (GFP) hepatocytes (CellTracker) 10kDa dextran
500 µm
C
Physiological Media - Day 6 (z-slices)
HUVECs (GFP) hepatocytes (CellTracker) 10kDa dextran
z = 39µm
z = 54µm
z = 69µm
z = 84µm
z = 99µm

**SI Figure 4: Vascularized liver devices are fully perfusable, with vessel structures surrounding and penetrating through hepatic spheroids.**

A-B: Maximum intensity projection images of devices show that perfusable vessels span the entire length of the device's central hydrogel compartment, under both physiological (A) and insulin resistance ("IR") (B) culture conditions. Vascular networks are formed from GFP-HUVECs (green) and lung fibroblasts (unlabeled), hepatocytes are labeled with CellTracker Deep Red (magenta), and perfusion is demonstrated using 10kDa dextran (blue).

C: Individual z-slices (spanning 60 µm in total height) show vessel structures (GFP-HUVECs, green) both wrap around and extend through a hepatic spheroid (hepatocytes, magenta). Bottom row of images shows that these vessel structures are perfusable (10kDa dextran, blue). Scale bars = 100 µm.

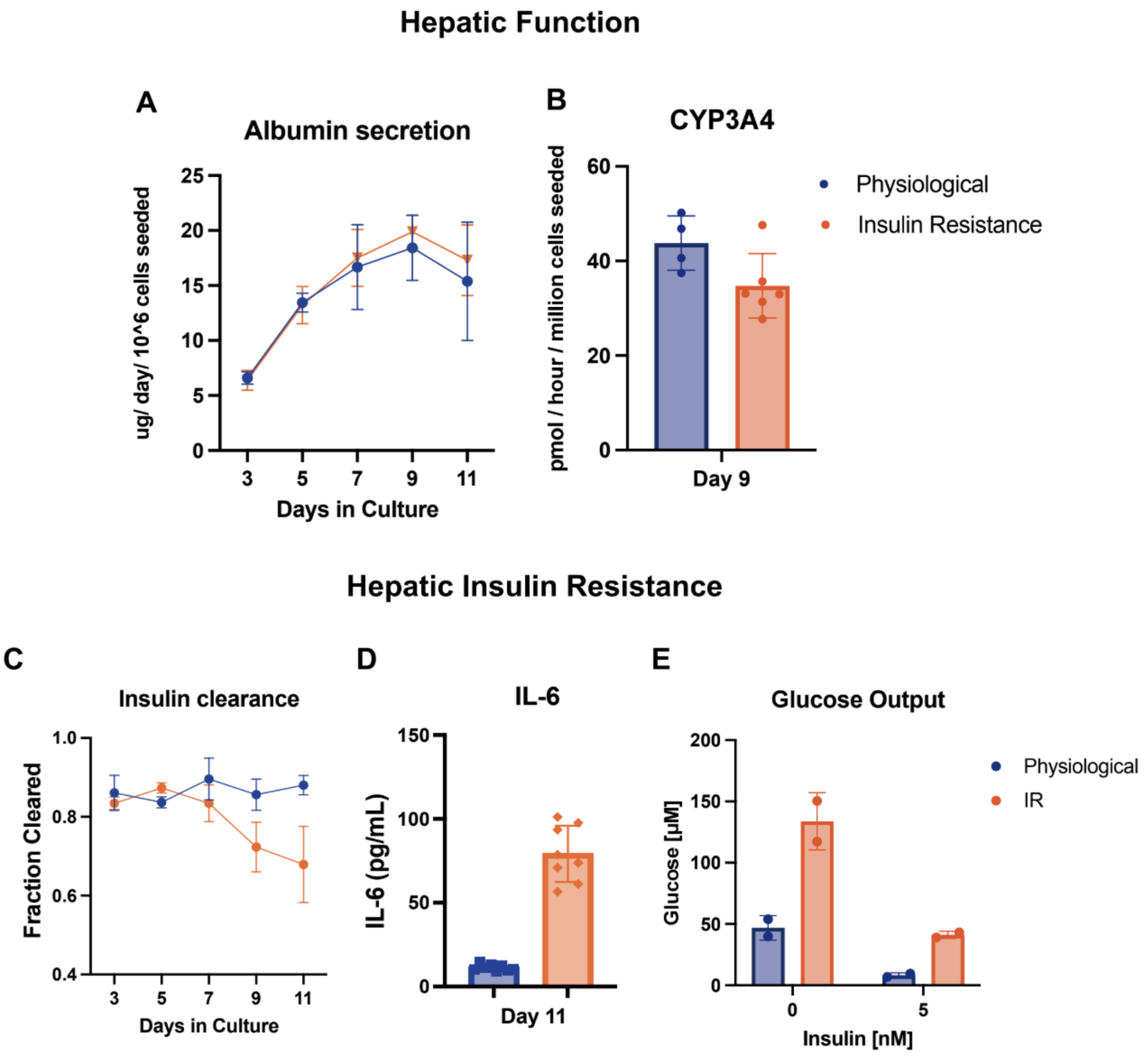


**SI Figure 5: Spheroid-only dataset demonstrates multicellular hepatic spheroids alone are functional and develop a disease phenotype after several days in culture in insulin resistance (IR) medium.**

A: Multicellular hepatic spheroids secrete albumin at comparable rates in both media conditions.

B: Spheroid CYP3A4 activity rate measured on day 9 in culture, normalized per million hepatocytes seeded.

C: Insulin clearance diminishes over time in IR samples, measured via ELISA every 48h.

D: Increased IL-6 secretion assessed on day 11 from spheroids cultured in IR conditions.

E: Glucose output is elevated over 24h from day 12 to day 13 in IR conditions, both with and without insulin stimulation.

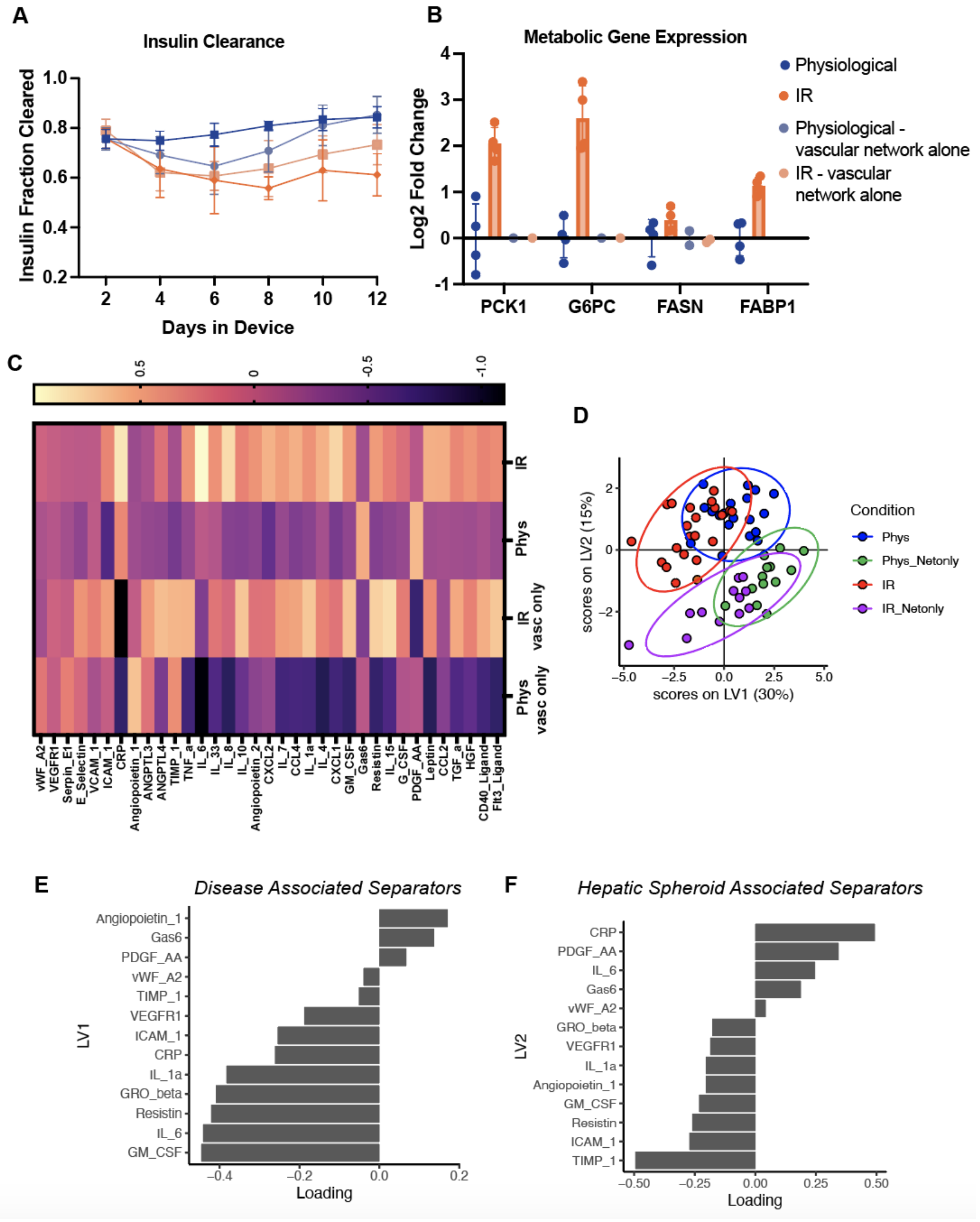

A
Insulin Clearance
Insulin Fraction Cleared
Days in Device
B
Metabolic Gene Expression
Log2 Fold Change
PCK1
G6PC
FASN
FABP1
Physiological
IR
Physiological - vascular network alone
IR - vascular network alone
C
IR
Phys
IR vasc only
Phys vasc only
vWF_A2
VEGFR1
Serpin_E1
E_Selectin
VCAM_1
ICAM_1
CRP
Angiopoietin_1
ANGPTL3
ANGPTL4
TIMP_1
TNF_a
IL_6
IL_33
IL_8
IL_10
Angiopoietin_2
CXCL2
IL_7
CCL4
IL_1a
IL_4
CXCL1
GM_CSF
Gas6
Resistin
IL_15
G_CSF
PDGF_AA
Leptin
CCL2
TGF_a
HGF
CD40_Ligand
Flt3_Ligand
D
scores on LV2 (15%)
scores on LV1 (30%)
Condition
Phys
Phys_Netonly
IR
IR_Netonly
E
Disease Associated Separators
LV1
Loading
F
Hepatic Spheroid Associated Separators
LV2
Loading
GRO_beta

**SI Figure 6: Vascular network alone devices show that vascular networks contribute to disease phenotype observations and state.**

A: Insulin clearance measurements comparing liver MPS with spheroid-free control devices show insulin is also cleared by vascular networks alone, and insulin clearance increases at approximately day 6 when vasculature becomes perfusable.

B: Gene expression of metabolic-associated genes measured via qPCR. PCK1 and G6PC were not detected in network alone controls.

C: Heatmap of analytes measured via Luminex comparing MPS with and without spheroids for each media condition.

D: Partial least squares discriminant analysis (PLS-DA) model built on Luminex data with feature selection shows separation between liver MPS (blue vs. red) and network alone controls (green vs. purple) as well as between media conditions within each of these groups.

E-F: Loadings on LV1 and LV2 identify analytes driving these separations identified in the PLS-DA model.

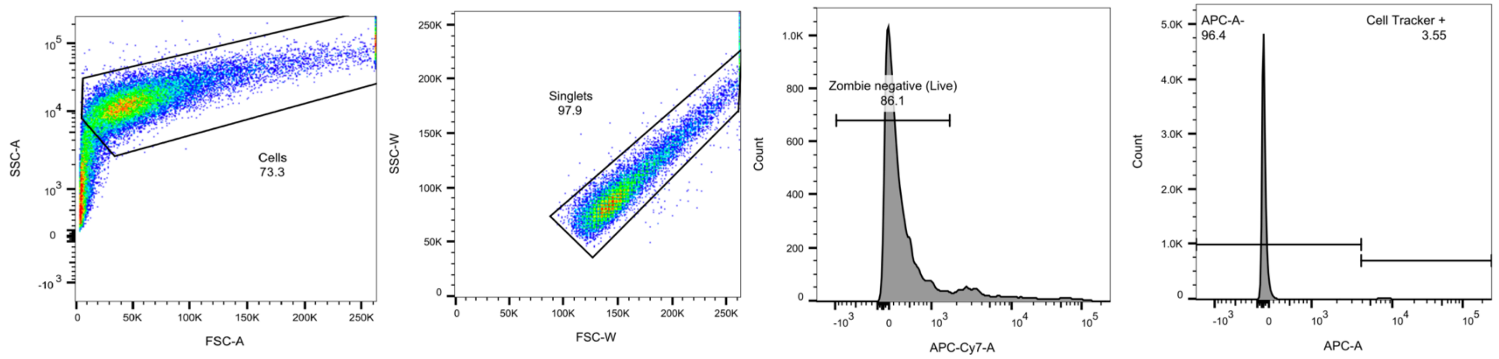


**SI Figure 7: Flow cytometry gating scheme for quantification of CellTracker+ monocytes.** Sequential gating for cell populations included identification of singlets by FSC-W vs. FSC-H and live cells as the Zombie-negative population. Monocytes were identified using APC filter settings to gate positive cells, which were stained with CellTracker Deep Red during initial seeding.

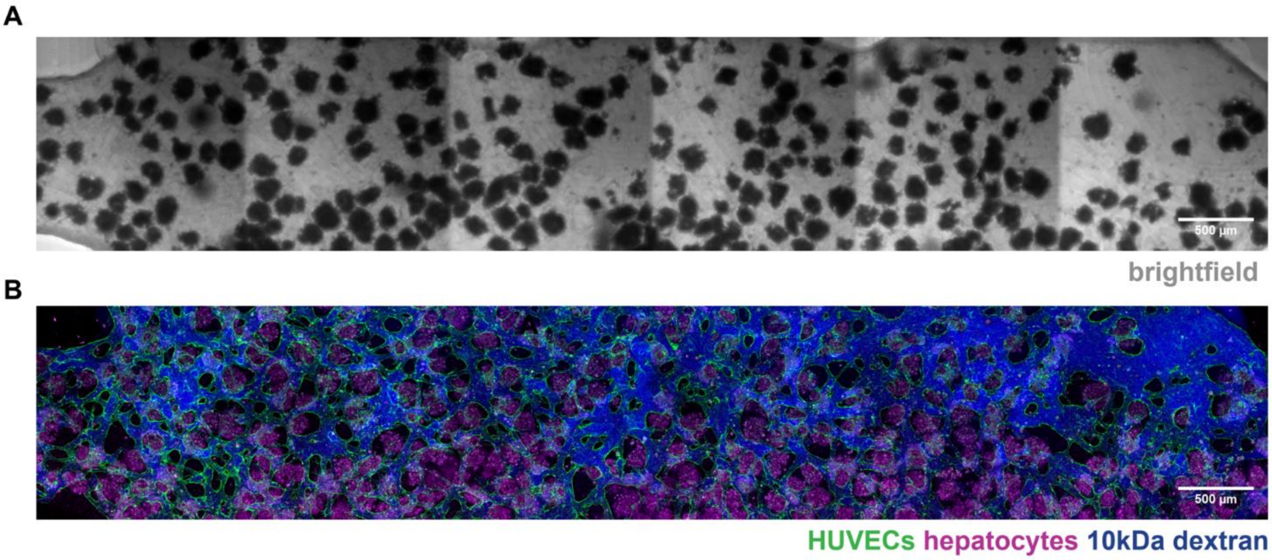


**SI Figure 8: Changing ratio of spheroids to cells that comprise the surrounding vascular networks is an avenue to improve physiological relevance.**
A: Maximum intensity projection brightfield image of a smaller vascularized liver device (hydrogel compartment dimensions are 1.8mm wide x 7mm long x 0.4 mm tall), taken on Day 7 with physiological media. In this smaller device, HUVECs and NHLFs were seeded at the same ratio as in the standard microfluidic device (8M/mL and 1M/mL, respectively), but the gel channel is 1/3 of the volume. The number of spheroids was kept constant (~200 spheroids per device), resulting in a higher ratio of spheroids to vascular network cells.
B: Maximum intensity projection image of the same device as (A). HUVECs are labeled with UEA-I rhodamine (false-colored green), hepatocytes are visualized via autofluorescence with 488 nm excitation (false-colored magenta), and perfusion is shown with 10kDa dextran (blue). Scale bars = 500 µm.

**Supplemental Videos**
Available at the following link: https://tinyurl.com/Tevonian-etal-SI-Videos

**Video 1: Widefield time-lapse video of vascular network formation in liver MPS devices over 12 days.** GFP-HUVECs self-assemble into vascular networks over time, connecting with RFP-HUVECs that migrate out from hepatic spheroids. Overlay of the brightfield channel (right side) shows spheroids. One image taken per day, scale bars = 250 µm.

**Video 2: Confocal z-stack video showing 3D interactions between perfusable vessels and spheroids.** After 6 days of device culture in physiological media, GFP-HUVECs (green) have formed vascular networks that wrap around and penetrate into hepatic spheroids (CellTracker Deep Red, magenta), with perfusion demonstrated using 10kDa dextran (blue). Each frame represents a 3µm slice.

**Video 3: Confocal maximum-intensity projection time-lapse video of CD14+ monocytes in devices over a 24-hour period.** CD14+ monocytes labeled with CellTracker Red (false-colored yellow) were added to GFP-HUVEC vascular networks (green) on Day 8 of device culture and imaged over 24 hours, with one z-stack image being taken every hour. Hepatocytes are labeled with CellTracker Deep Red (magenta).
